# Feasibility of Remotely Supervised Transcranial Direct Current Stimulation and Cognitive Remediation: A Systematic Review

**DOI:** 10.1101/768879

**Authors:** Nicole Gough, Lea Brkan, Ponnusamy Subramaniam, Lina Chiuccariello, Alessandra De Petrillo, Benoit H. Mulsant, Christopher R. Bowie, Tarek K. Rajji

**Author notes:** Centre for Addiction and Mental Health, 80 Workman Way, Room 6312, Toronto, ON Canada M6J 1H4. **Authors Statement:** NG, BL, SP, ADP, CRB, and TKR contributed to conception, design, analysis and interpretation of data. NG, BL, SP, and ADP contributed to data acquisition. All authors contributed to the drafting and critically reviewing the manuscript and approved the final version.

## Abstract

With technological advancements and an aging population, there is growing interest in delivering interventions at home. Transcranial Direct Current Stimulation (tDCS) and Cognitive Remediation (CR) have been widely studied, but mainly in laboratories or hospitals. Thus, the objectives of this review are to examine feasibility and the interventions components to support the domiciliary administration of tDCS and CR. We performed a systematic search of electronic databases, websites and reference lists of included articles from the first date available until October 31, 2018. Articles included had to meet the following criteria: original work published in English using human subjects, majority of tDCS or CR intervention administered remotely. A total of 39 studies were identified (16 tDCS, 23 CR/cognitive training, 5 using both tDCS & cognitive training). Four studies were single case studies and two were multiple case studies. The remaining 33 studies had a range of 9 – 135 participants. Five tDCS and nine CR/cognitive training studies were double blind randomized controlled trials. Most studies focused on schizophrenia (8/39) and multiple sclerosis (8/39). Literature examined suggests the feasibility of delivering tDCS or CR remotely with the support of information and communication technologies.

## INTRODUCTION

Currently, 47 million people worldwide suffer from dementia, with nearly 10 million new cases each year, making it the 7th global leading cause of death [1]. Alzheimer’s Dementia (AD) represents a growing health concern and contributing to 60–70% of dementia cases worldwide [2, 3]. Given the growing prevalence rate of AD, preventative interventions and treatments that target individuals on a population level are crucial.

There are limited effective treatments available for AD, highlighting the necessity for preventative options. Recent research on preventative measures has focused on interventions that target brain neuroplasticity and cognitive reserve [4] due to observed maladaptive neuroplastic changes in various neuropsychiatric diseases [5]. Such changes are also visible in AD, whereby deficits in cognition, may be related to disruptions in the connections among neurons and neuronal networks [5]. Therefore, inhibiting these pathological changes and enhancing neuroplasticity may be beneficial for preventing or delaying the onset of AD and enhancing cognition [5]. Two interventions that have the potential of enhancing neuroplasticity and can be delivered remotely, offering a scalable preventative effect, are transcranial direct current stimulation (tDCS) and cognitive remediation (CR) [4, 6].

tDCS is a non-invasive brain stimulation method that can be safely administered to awake outpatients and works by delivering a low intensity electrical current (e.g., 2 mA) to either increase cortical excitability with an anodal electrode, or to suppress cortical excitability with a cathodal electrode [7]. tDCS has the potential to reduce symptoms of cognitive decline and enhance cognition and rehabilitation in neuropsychiatric diseases [5, 8], including mild AD [9–15], through the modulation of neuronal activity and neuroplasticity [5, 7]. Unlike other non-invasive brain stimulation devices, tDCS equipment is readily transportable, making it a viable population-level intervention, and an option for patients with AD who need remote assistance.

CR programs also offer a way of improving neurocognitive abilities by inducing functional changes within the brain [4]. In CR, patients engage in computerized cognitive exercises and, with the help of a therapist, are encouraged to utilize their metacognition in order to identify and modify their problem-solving techniques [16]. CR has been shown to improve cognition in schizophrenia [15], bipolar disorder [12, 13], alcohol dependence [14], and major depression [4, 8, 17]. CR training programs are available online and therefore can be easily accessed from home.

Although a number of studies have examined the tolerability and efficacy of home-based, remotely-supervised tDCS and cognitive training in order to determine viability [18–20], no study to date has examined the combination of tDCS and CR delivered remotely as preventative measures for dementia. The current review intends to summarize existing literature on remote delivery of tDCS and CR/CT. Insight into the current research findings will allow for future determination of the potential usefulness of these two techniques to act as preventative treatment options for dementia on a population level.

## METHODS

### Selection strategy

PubMed, Ovid, PsycINFO and CINAHL databases were searched focusing on studies from the first date available to October, 31 2018 for tDCS and CR/CT at-home studies. The literature search was divided into 2 categories:

#### tDCS at-home studies

Titles and abstracts for the following Medical Subject Headings (MesH) terms and keywords: (transcranial direct current stimulation or tDCS) in combination with “at-home” OR “home-based” OR “remotely supervised” OR “home treatment” OR “telemedicine” OR “self-administered” were searched.

#### Cognitive Training and Cognitive Remediation at-home studies

Titles and abstracts for the following MesH terms and keywords were searched: (cognitive remediation) in combination with “at-home” OR “home-based” OR “remotely supervised” OR “home treatment” OR “telemedicine” OR “self-administered.”

### Selection Criteria

#### tDCS at-home studies

The following inclusion criteria were applied: (1) articles published in English; (2) original research with human participants, (3) home-based intervention. Excluded papers were: (1) articles reporting tDCS data from research settings, such as laboratories, hospitals, clinics, and research centers; (2) review, guideline and protocol papers without reporting original research; (3) articles that used animals as study subjects, (4) articles reporting do-it-yourself (DIY) tDCS use.

#### Cognitive Training and Cognitive Remediation at-home studies

The following inclusion criteria were applied: (1) articles published in English; (2) original research with human participants; (3) home-based interventions. Excluded papers were: (1) articles reporting CR/cognitive training data from research settings such as laboratories, nursing homes, hospitals, clinics, and research centers, (2) review, guideline and protocol papers without reporting original research.

### Data Extraction/Collection

Data extraction is illustrated in Figures 1 and 2. After filtering for inclusion and exclusion criteria and eliminating duplicates, 11 home-based tDCS papers, 23 home-based cognitive training (CT) or CR papers, and 5 studies that discussed both home-based CT and tDCS were included from this systematic multiple database search in the current review.

**Figure 1.**
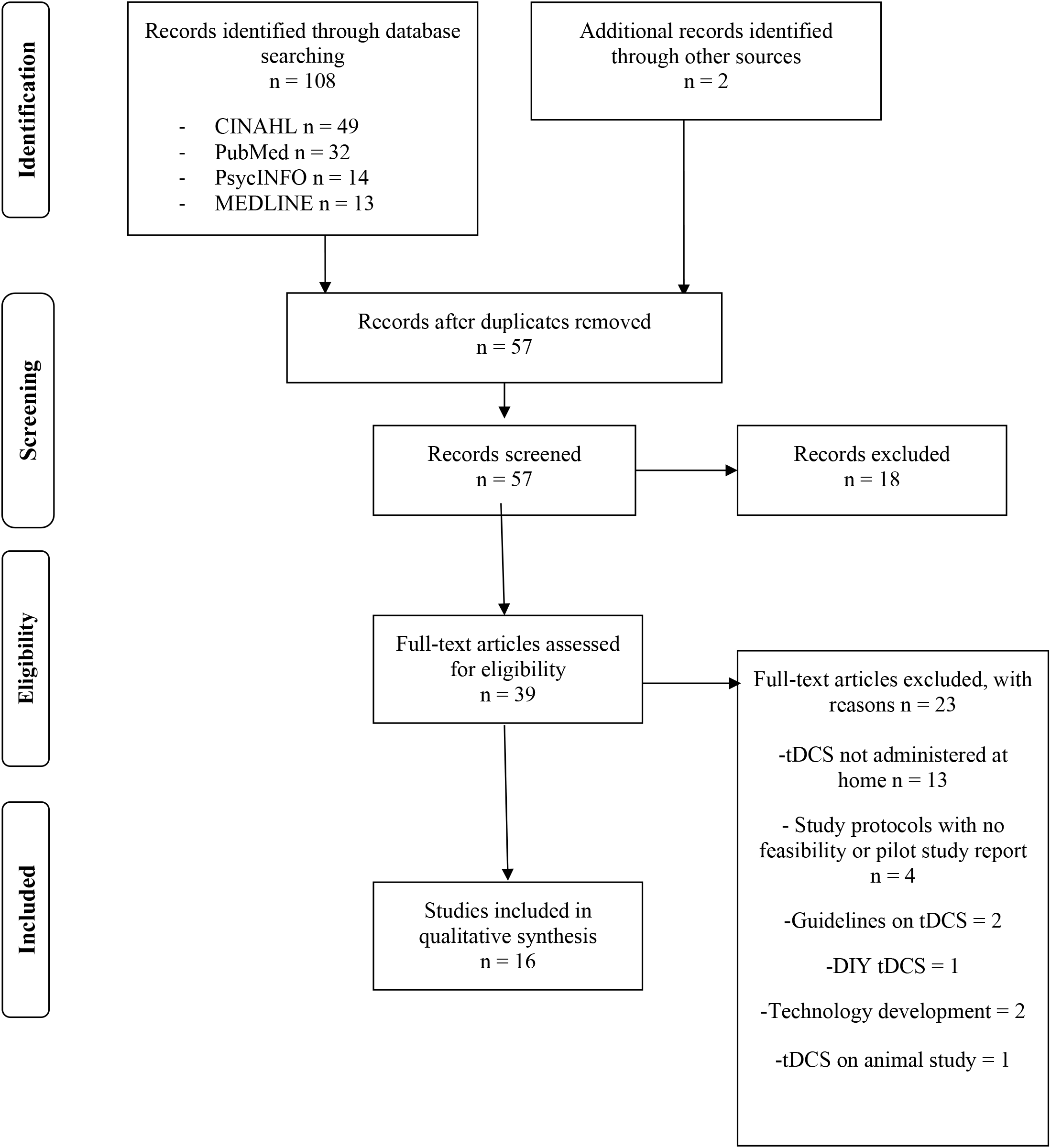
Preferred Reporting Items for Systematic Reviews and Meta-Analyses (PRISMA) Flow Diagram for tDCS At-Home Studies

**Figure 2.**
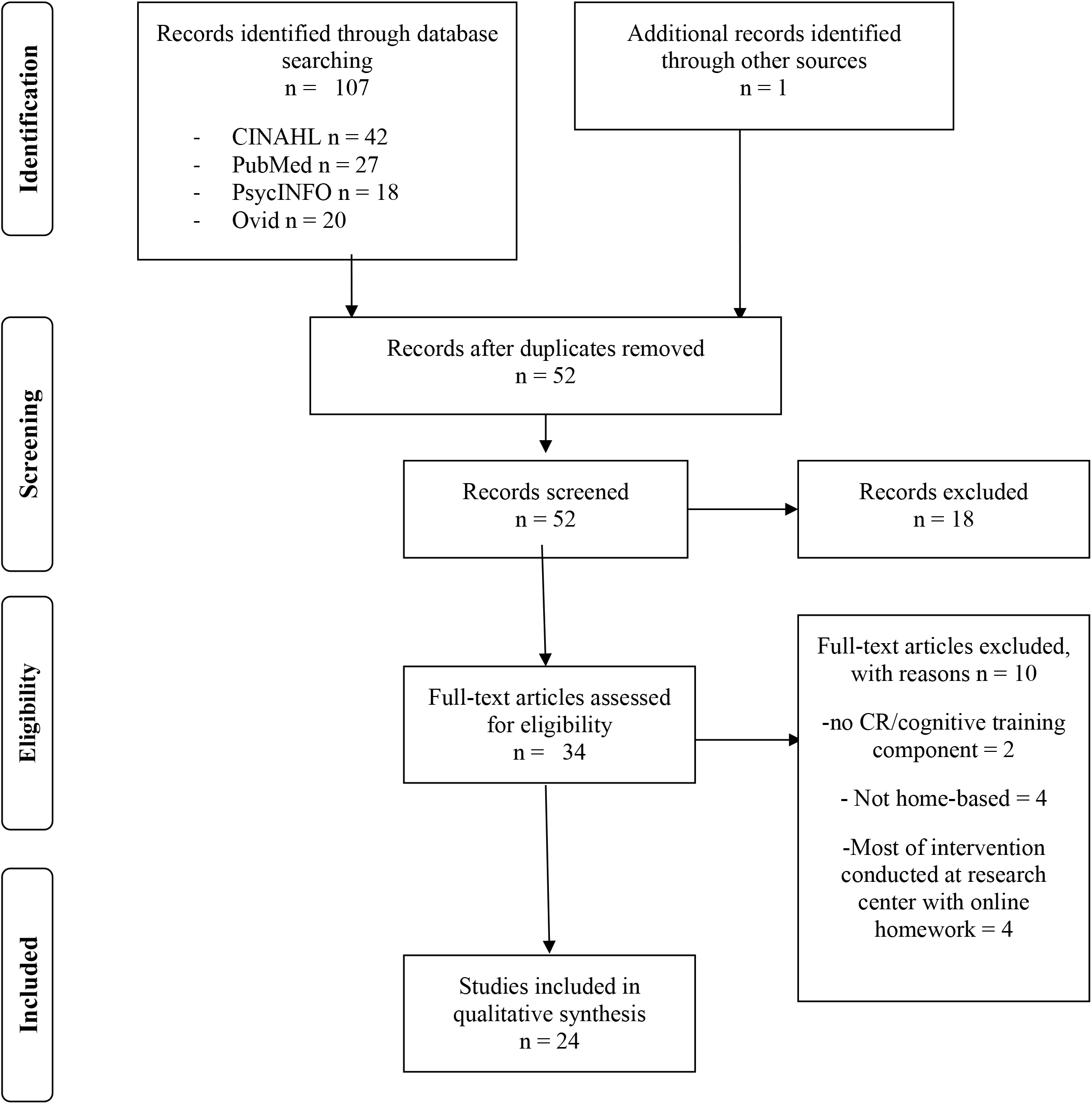
Preferred Reporting Items for Systematic Reviews and Meta-Analyses (PRISMA) Flow Diagram for Cognitive Remediation and Cognitive Training At-Home Studies

## RESULTS

A total of 39 publications met the inclusion criteria for this review. Out of the 39 identified publications, two studies were single case studies and two were multiple case studies, one involving six participants and the other involving four participants. The remaining 33 published articles reported on a sample size range of 9-135 participants. Studies were published between the years of 1995-2018, and targeted multiple diagnoses including but not limited to: multiple sclerosis [2, 19, 20, 26–28, 45], Parkinson’s disease [2, 30, 43], brain injury [44, 52], and schizophrenia [29, 37, 46, 48, 49, 51, 55]. Only three papers [21–23] and one case study [24] were published with AD as the target population, with an additional study targeting mild vascular dementia [25]. Studies examined in this systematic search included both males and females with a noted age range of 7.77 (standard deviation: 1.62) to 94 years of age.

### Remote tDCS Research Published to Date

There were 16 studies total that met the inclusion criteria. Studies targeted a range of symptoms and disorders, but most research focused on multiple sclerosis [2, 20, 26–28], schizophrenia [29], Parkinson’s disease [2, 30] and dementia [24, 25].

### Study Design

Of the 16 tDCS studies reviewed, five were double blind randomized controlled trials [2, 27, 31-33], two were single blind randomized sham controlled trials [25, 34], two were pilot studies [20, 28], two featured a randomized double-blind cross-over design [18, 35], two were open label studies [27, 30], one was a multiple case study with four participants [36] and two were single case studies [24, 29].

### Participants

There was high variability in the sample size and age of participants across studies. The ages of participants ranged from 17 to 86. The number of participants in each study ranged from 1 to 45, with an average of 19.63 (SD 12.02) participants.

### Administration Parameters

tDCS stimulation was administered for 20 minutes per session in all studies except three [24, 29, 33]. The current used for the stimulation ranged between 1 to 3mA, with 12 studies using either 1.5mA or 2mA.

Participants either self-administered tDCS (n=5/16 studies) [33], administered tDCS with the help of an aide or study partner (n=6/16 studies) [18, 24, 25, 29, 31, 35, 36], or could choose what they preferred [36]. Five studies encouraged self-administration, but used a proxy or caregiver if further assistance was required [20, 26–28, 30].

The most common electrode placement was the dorsolateral prefrontal cortex setup (DLPFC), where the anodal electrode was placed over the left DLPFC, and the cathodal electrode over the right DLPFC or temporal lobe (n=12/16 studies) [2, 21, 26–30, 32, 33, 35, 36]. Furthermore, the majority of studies reviewed utilized tDCS head gear for simple and consistent electrode placement each session (n=11/16 studies) [2, 20, 26–30, 32–34, 36].

The number of sessions administered varied across studies with participants receiving an average of 15.17 (SD = 12.36) tDCS sessions, ranging between 4 to 60 sessions for studies lasting less than four months. Within longer running studies, one study involved 8 months of daily tDCS sessions [24], and another study followed a participant for three years receiving 1-2 daily treatments, with tDCS sessions still ongoing at the time of publication [29].

### Training and Safety Measures

Although the majority of studies employed tDCS training at the baseline testing visit or during the first session at the research center (n=12/16 studies) [2, 18, 20, 26–28, 30–33, 35, 36], the length, intensity and nature of training varied across research. In most cases training sessions consisted of an instructional video, hands-on tDCS coaching and monitoring by a study technician [2, 26, 28, 30, 33, 35, 36], troubleshooting techniques [2, 28, 33, 35, 36], as well as an assessment of the participant’s ability to replicate the procedures competently at home [2, 32–36]. Training also incorporated safety assessments, including tolerability testing at the initial clinic visit [2, 20, 26-28, 30, 33, 36].

Most studies implemented a number of safety measures in order to ensure safe and controlled use of the tDCS device. This included tDCS devices that were programmed to allow for a minimum of 12 hours between sessions [33], and devices that released a single session of stimulation after receiving a one-time unlock code [2, 20, 26–28, 30, 33, 36]. Additional safety measures included machines with password protected settings [34], anode and cathode sponges that had opposite male and female connections [34], as well as tDCS machines that measured and displayed contact quality prior to the start of stimulation [2, 20, 28, 30, 33, 35, 36].

### Fidelity Monitoring

All studies (n=16 studies) used some form of fidelity monitoring including real-time monitoring by means of video-conferencing (n=8/16 studies) [2, 20, 26–28, 30, 33, 36], remote control software (n=7/16 studies) [2, 20, 26–28, 30, 36], daily online check-ins (n=1/16 studies) [34], treatment diaries (n=4/16 studies) [18, 32, 33, 35], webcam or Facetime sessions (n=1/16 studies) [34] and weekly home visits (n=1/16) [35].

### Effectiveness of tDCS Treatment

Overall, all studies reviewed reported significant improvement in physical or mental symptoms, such as a reduction in fatigue [20, 27] and pain [18, 20], improved pain management [35], improved anxiety and dizziness ratings [34], improvements in affect [20], motor function [30], rocking perception, [34], psychosocial functioning [29] and moderate improvements in consciousness [35]. Further significant improvements were noted in areas of cognition, including processing speed [20, 25], visual recall [25], attention and response variability composites [26] as well as stabilization of cognitive decline for AD patients, with some improvement in memory function also noted [24].

### Feasibility of Remote tDCS

Many studies supported the feasibility of remotely supervised tDCS [2, 20, 30, 33, 35, 36] and found it to be well tolerated [18, 20, 30, 33, 35], easy to use and safe [32, 33]. High compliance rates (80% or higher) were noted in nine studies [19, 30, 33, 35, 37–41]. Only one study noted generally poor compliance rates and a high dropout rate [42], and two other studies noted participants training less than desired [43], or not always following the protocol [44]. The remaining four studies did not discuss compliance rates.

Five studies addressed the combination of remotely delivered tDCS and cognitive training [2, 20, 26, 27, 30]. No additional training or supervision by study staff was discussed with the incorporation of cognitive training into the study protocol.

### Remote Cognitive Remediation and Cognitive Training Research Published to Date

There were 24 studies total that met the inclusion criteria. There was wide variability in the populations targeted including, but not limited to, schizophrenia [37, 46, 48, 49, 51, 55] and individuals who were at risk of developing psychosis [39], multiple sclerosis [2, 19, 45], dementia [21–23], Parkinson’s disease [2, 43], and brain injury [44, 52].

### Study Design

Among the 24 studies reviewed, there were nine double blind, randomized controlled trials [2, 19, 21, 22, 39, 40, 42, 45, 46], one randomized waitlist controlled trial study [47], one multicenter randomized control trial [23], three pilot studies [41, 48, 49], one single group study [43], one within subject study [50], one controlled experimental design [51], one follow-up single group design study [52], one case control study [53], one longitudinal within-group design study [38], one feasibility study [37], one multiple case study [54], and two single case studies [44, 55].

### Participants

The age range and sample size varied, with participants ranging in age from 7 to 75. The number of participants in each study ranged from 1 to 135, with an average of number of 41.1 (SD =35.55) participants.

### Administration Parameters

There was high variability in the administration of CR and CT. The duration of engagement, with one exception [44], ranged from 20 minutes per day to two hours per day, with a frequency of two to seven days per week.

There was some variability in the type of CR or CT delivered, as well as the mode of delivery. Of the 24 studies examined, six studies utilized principles of cognitive remediation [22, 23, 38, 42, 48, 52–54], whereas the other 18 studies utilized cognitive training exclusively. Participants completed tasks individually (n=14/24 studies) [2, 19, 38, 39, 42, 43, 45, 46, 48–53], or with a partner (n=10/24 studies) [21-23, 37, 40, 41, 44, 47, 54, 55]. Studies used printed materials (n=7/24 studies) [21-23, 42, 44, 54, 55] and internet-based and/or computer-based programs (n=18/24 studies) [2, 19, 38, 39, 42, 43, 45, 46, 48–53] to deliver the intervention to participants. Only five studies did not use any form of computerized program [21-23, 52, 55]. Software also varied quite extensively in studies that utilized a computer. Posit Science was used in six different studies [19, 37, 39, 45, 46, 50], however, the program varied in its usage and utility.

A common factor across studies was utilizing programs that adjusted in difficulty level based on participant performance (n=21/24 studies) [2, 19, 21, 22, 37–41, 43–47, 49–55].

### Training and Guidance

Participant training ranged in duration and intensity and consisted of no training and self-guided treatment (n=4/24) [38, 41, 45, 52], having one session of training in a group or alone (n=5/24) [2, 37, 40, 42, 47], weekly training [44], observational training [21, 22, 53] or training on an as-needed basis [46]. Training generally involved printed instruction sheets and recommended strategies for daily activities, which were explained prior to starting the intervention [54], education surrounding the human brain, cognition and how cognition affects daily functioning [37, 42], educational handbooks, worksheets and information about compensatory strategies [42], as well as the protocol for the computer software [50]. Only three studies discussed initial training, whereby training was conducted at the study center during the first session [42, 48] for two hours [37].

### Fidelity Monitoring

Most research provided ongoing monitoring and support by a study technician (n=20/24 studies) [2, 19, 38, 39, 42, 43, 45, 46, 48–53]. Software programs that allowed for monitoring/real time feedback were utilized in most studies (n=14/24) [2, 20, 38–40, 42–44, 45, 46, 47, 49, 50, 52, 54]. Check-ins were also quite common (n=16/24 studies) [2, 19, 38, 39, 42, 43, 45, 46, 48–53], and were completed primarily through phone or email at least once a week (n=7/24) [19, 37, 39, 40, 45, 49, 51]. Other studies used video conferencing multiple times a week [38], or had study staff meet with participants once a week either at home [54, 55] or in a study clinic [54]. Additional tools included: instructional manuals [23, 24, 39, 54, 55], participant logs [50], checklists [54], and training schedules [45, 47, 55].

### Effectiveness of Remote Delivery of Cognitive Remediation and Cognitive Training

Overall, all remote cognitive remediation and cognitive training studies reviewed showed a positive effect in most of the parameters measured following the remote intervention. Specifically, significant cognitive improvements were found for individuals with MS [19, 45], chronic fatigue syndrome [53], 22q11 Deletion Syndrome [38], schizophrenia [34–36, 55], HIV [50], TBI [44], acquired brain injury [52], and AD [21]. Areas of cognition that were improved following these interventions include, but are not limited to, global cognition (n=9/24) [19, 21, 37, 39, 43, 45, 46, 51, 55], processing speed (n=3/24) [49, 50, 53], working memory (n=3/24) [22, 44, 53], visual recognition (n=1/24) [44], verbal memory (n=2/24) [46, 51], word fluency (n=3/24) [21, 22, 44], and executive functioning skills (n=5/24) [22, 46, 47, 51, 53]. Only one study did not find improvements in cognition, however, they did find statistically significant improvements within activities of daily living performance for stroke patients with cognitive impairments [54]. Many studies found that improvements in areas of cognition persisted at least 3-9 months post-intervention (n=5/24) [21, 40, 41, 43, 51]. Improvements in cognition were also found to be positively correlated with quality of life (n=3/24) [42, 44, 51].

### Feasibility of Remote Cognitive Training and Cognitive Remediation

Most research reported remote, home-based cognitive remediation and cognitive training programs to be feasible, useful and well accepted for people with MS [19, 45], schizophrenia [48, 55], epilepsy [42], and those with brain damage [44, 52]. Participants indicated that the training was beneficial, convenient [42], and enjoyable [42, 48], and supported the benefit of remote CR using spousal caregivers for people with dementia [22]. Only one study noted that some participants felt the cognitive training instructions were confusing [50].

## DISCUSSION

The current literature suggests that both remotely administered tDCS and CR are feasible and effective in targeting a number of cognitive functions in various patient populations. Proper training in equipment use and regular monitoring of procedures appear crucial for study compliance and feasibility. The literature suggests that implementing ideal components of both CR and tDCS administration should be studied as a viable preventative treatment option for AD.

### tDCS

Based on the literature reviewed, the recommendations for future trials targeting cognitive decline in the AD population would include ensuring proper training and regular monitoring. This means dedicating adequate training time at the initial clinic visit or arranging in-home training in order to ensure hands-on coaching around tDCS application and proper use of the device. Explanation and demonstration of troubleshooting techniques could be helpful to include as part of the training procedure, including rectification techniques to manage pain and discomfort.

Considering the potential difficulty in tDCS self-administration with a predominantly elderly population, machine safety features and a customized headset could be useful. If a headset is not used, study partners or proxies can be utilized for tDCS application, with an anode and cathode that are clearly distinguishable to avoid interchangeability.

Video-monitoring as well as online and in-person check-ins could be beneficial ways for ensuring fidelity monitoring for a protocol targeting cognitive decline in the AD population. It is recommended that participants in future studies be overseen once every week, or every two weeks, through video conferencing or home visits by study staff. If tDCS stimulation is combined with an online cognitive training program, monitoring could also include remote control software. In addition, it is recommended that participants keep a log of sessions completed, either digitally or manually, in order to ensure protocol adherence.

The effectiveness of tDCS was dose-dependent, where more sessions (20 vs 10) and an increase in current (2.0mA vs 1.5mA) resulted in a greater reduction in fatigue [27, 29]. This suggests that longer treatment periods and higher stimulation intensity are of greater benefit where at least 20 tDCS sessions with a current no lower than 2.0mA [18–20, 56] may be optimal. Future trials might want to consider these settings to ensure maximum benefit.

### Cognitive Remediation

Ease of use may be a necessary factor to take into account for future trials in the AD population. Considering these patients already suffer from some cognitive decline and may be less experienced with computer software, a simpler computer program preinstalled by study staff and a longer training period might be preferred for tolerability. Additionally, having study staff engage in regular monitoring (i.e. regular phone calls, home visits or video conferencing) and using programs that ensure optimal cognitive challenge appear to be important components to consider for effectiveness and adherence.

Future trials may also want to consider utilizing exercises that target cognitive domains typically affected by AD related cognitive decline, such as memory, executive function, processing speed, attention, as well as reasoning and problem solving.

Moreover, continuously challenging cognition by finding an individualized optimal level of difficulty within the CR program used is an important aspect to consider for future trials targeting AD related cognitive decline. With the exception of four studies [23, 42, 48, 52], all research reviewed had cognitive exercises that adapted in difficulty level based on the participant’s performance. This self-adjusting feature may promote continued participant engagement and eliminate frustration brought about by seemingly unachievable difficulty parameters within the exercises.

Strategy contemplation [38, 42, 52] and cognitive transfer [22, 42, 52, 54] are also important components to consider within the study design, whereby participants are asked to identify skills and strategies utilized in the exercises and apply them to real-world situations. This addition encourages cognitive activation and the utilization of adaptive problem-solving strategies in daily scenarios. To further cognitive transfer, some CR interventions encourage participants to seek out cognitively challenging activities in everyday life, and engage in activities that are cognitively stimulating outside of the program, however, this was not clearly visible in any of the studies reviewed. These add-on techniques could be particularly helpful in slowing cognitive decline and facilitating improvement in every day functioning and daily living within the AD population.

### Limitations

A major limitation of this systematic review is that out of 39 studies reviewed, only four presented AD as the disease targeted. More specifically, within the cluster of at-home tDCS literature reviewed, only one case study used the intervention as a means to slow cognitive decline in an individual with early onset AD [24]. Similarly, within the group of at-home CR articles reviewed, one RCT targeted patients with AD [22], and one targeted individuals with Mild Cognitive Impairment (MCI) and AD [23]. A third article targeting AD utilized cognitive training rather than cognitive remediation [21]. This makes it difficult to make inferences and recommendations based on population specific difficulties encountered in past studies that could be improved upon in future studies.

Only six studies utilized discussion guided cognitive remediation (rather than cognitive training) as the observed intervention, with aspects of strategy awareness [55] and discussion [23, 38, 42, 52] visible in five out of six CR studies reviewed, and the addition of cognitive transfer visible in 4 out of 6 [22, 42, 52, 54]. As a result, it is difficult to evaluate the effectiveness and value of these add-on techniques within a trial targeting patients at-risk for developing AD. Furthermore, as different cognitive measures were used in each of the studies, it is difficult to determine if one method was more effective over another.

## CONCLUSIONS

With millions of individuals being diagnosed with AD worldwide each year, and the accompanying maladaptive neuroplastic changes leading to worsening cognition, CR and tDCS are promising preventive interventions and treatments that can target individuals on a population level. Both CR and tDCS are effective interventions for improving cognition [4, 8–15, 17] and have the benefit of being portable.

CR enhances frontal lobe activation and neuroplasticity [57] and it has been shown to improve cognition in depression [4, 8, 17]. CR’s effectiveness relates to its inclusion of performance adapting software as well as strategy-based learning and bridging discussions, which typically occur in a group environment. However, with current technological advances, these components of CR can be achieved remotely.

tDCS modulates neuronal activity and enhances neuroplasticity [7] and has been shown to improve cognition in mild AD [9–11]. In other studies, participants must be present at a treatment centre in order to receive CR, tDCS or both from trained study staff. However, this can be costly and laborious for participants, and has the potential of being unfeasible for individuals with restricted mobility, vocational obligations, and lengthy travel times.

Future research may like to further consider additional measures that were not fully assessed throughout the literature examined, such as baseline ratings of reward (which may be positively associated with cognitive gains) and inclusion of a partner or caregiver. Investigating the feasibility of remotely delivering these interventions with other cohorts, such as individuals with mild AD, or those at risk of developing AD, who could benefit from at-home tDCS, CR or both would also be important considerations for prospective trials.

**Table 1:** Summary Characteristics of Studies on Remotely-Delivered tDCS.

| Authors (Year) | Type of Study | Disease | N | Age | tDCS Current (mA) | Number of tDCS Sessions | Duration of tDCS Stimulation | Electrode Placement | Results |
| --- | --- | --- | --- | --- | --- | --- | --- | --- | --- |
| Agarwal et al. (2018) | Open label study | Parkinson Disease (PD) | 16 Enrolled<br>10 in final analysis | 67.6 ± 5.9 | 2.0 | 10 | 20 Min | Bilateral DLPFC Montage (Left Anodal) | Significant improvement in motor symptoms. |
| Andrade (2013) | Single case study | Schizophrenia | 1 | 25 | Session 1-5 = 1.0. Session 6+, 2.0, then 3.0 | 1 or 2 sessions per day for 3 years | Sessions 1-5 = 20 Min. After session 5 = 30 Min | Anodal tDCS over left DLPFC and cathodal over left temporoparietal cortex. | Greater improvement in psychosocial functions. |
| Andre et al. (2016) | Single blind randomized sham controlled trial | Mild Vascular Dementia | 21 (13 active & 8 sham) | 78.6 (age range: 63-94) | 2.0 | 4 consecutive sessions | 20 Min | Anodal or sham over left DLPFC | Anodal stimulation showed meaningful improvement in visual recall and reaction times. |
| Bystad et al. (2017) | Single case study | Alzheimer's Disease | 1 | 60 | 2.0 | Daily for 8 consecutive months | 30 Min | Anodal over left temporal lobe (T3 in the 10/20 system) and reference electrode over right frontal lobe | Cognitive function was stabilized; improved immediate and delayed recall. |
| Carvalho et al. (2018) | Double blind randomized controlled trial | Healthy Subjects (HS) and Fibromyalgia (FM) | <u>HS</u>: 20 enrolled 19 in final analysis<br><u>FM</u>: 8 | <u>HS</u>: 26.31 ±4.89<br><u>FM</u>: 49.5 ±8.48 | 2.0 | <u>HS</u>: 10<br><u>FM</u>: 60 (5 session per week) | <u>HS</u>: 20 Min<br><u>FM</u>: 30 Min | <u>HS</u>: Anodal; Left primary motor cortex (M1) & Cathodal; contra-lateral supra-orbital area.<br><u>FM</u>: Left DLPFC | The findings suggest tDCS is feasible for home use with monitoring. |
| Cha et al. (2016) | Single blind randomized sham controlled trial | Mal Debarquement Syndrome | 24 (12 active 10 Sham & 1 open label) | 52.9 (12.2) | 1.0 | 20 (5 sessions per week) | 20 Min | Anodal placed over left DLPFC and cathodal over right DLPFC. | Active tDCS after rTMS improved rocking perception, anxiety and dizziness. |
| Charvet et al. (2017) | Double blind randomized controlled trial | Multiple Sclerosis | <u>Study 1</u> 15 Active; 20 Control<br><u>Study 2</u> 15 Active; 12 Sham | <u>Study 1</u> 52<br><u>Study 2</u> 44.2 | Study 1 = 1.5.<br>Study 2 = 2.0. | Study 1: 10<br>Study 2: 20 | 20 Min | Anodal was placed over the left dorsolateral prefrontal cortex (DLPFC) | tDCS has the potential to significantly reduce multiple sclerosis related fatigue. |
| Charvet et al. (2017) | Open label study | Multiple Sclerosis | 45<br>25 tDCS + CT;<br>20 CT Only | 51.96 (11.0) | 1.5 | 10 | 20 Min | Used “OLE” system, targeted DLPFC; Anodal on the left (F3), cathodal over right (F4). | Anodal stimulation at both sites improved complex attention and response variability composites compared to CT only group. |
| Hagenacker et al. (2014) | Randomized double-blind cross-over design | Trigeminal Neuralgia | 17 enrolled, 10 completed study | 63 (age range 49–82) | 1.0 | 14 consecutive daily sessions | 20 Min | Anodal tDCS over the primary motor cortex (M1). | Pain intensity significantly reduced. |
| Hyvarinen et al. (2016) | Double blind randomized controlled trial | Tinnitus | 35 (active tDCS = 23 & 12 sham) | 51 (15.4) | 2.0 | 10 consecutive sessions | 20 Min | Two different placements (1) Anodal over left temporal area & cathodal over frontal area; (2) Anodal & cathodal placed symmetrically bilaterally over frontal areas. | Overall improvement in tinnitus severity. |
| Kasschau et al. (2015) | Pilot study | Multiple Sclerosis (MS) | 20 (4 with proxy) | N/A | 1.5 | 10; over period of 2 weeks | 20 Min | Electrodes were placed in the bilateral dorsolateral prefrontal cortex (DLPC); Anode placed over left side. | Feasibility of remotely-supervised tDCS established for MS patients with Expanded Disability Status Scale (EDSS) of 6.0 or below OR 6.5 or above with proxy. |
| Kasschau et al. (2016) | Pilot study | Multiple Sclerosis | 20 (all active) | 51 (9.25) | 1.5 | 10 | 20 Min | DLPC; uniform bilateral dorsolateral prefrontal cortex (left anodal). | Anodal stimulation improved all symptoms measured; pain, fatigue, affect and cognitive processing speed. |
| Marten et al. (2018) | Randomized double-blind cross-over design | Minimally Conscious State (MCS) | 37 enrolled;27 Final analysis: 12 active/ sham 10 sham/ active | Age range: 17-75 | 2.0 | 20 sessions over period of 4 weeks | 20 Min | Anodal: over left DLPFC & cathode over right supraorbital region. | Moderate improvement in recovery of signs of consciousness.<br>-Only 17 patients completed remote tDCS. The rest completed tDCS from nursing home/rehab center. |
| Mortensen et al. (2015) | Double blind randomized controlled trial | Stroke-Patients with upper limb motor impairment following intracerebral hemorrhage | 15 (8 anodal tDCS, 7 sham) | 44 - 76 | 1.5 | 5 consecutive sessions | 20 Min | Anodal or sham over primary motor cortex (M1); anode placed on ipsilesional MI and cathode over contralesional supraorbital region. | Anodal tDCS + occupational therapy (OT) provide greater improvements compared to OT only. |
| Riggs et al. (2018) | Multiple case study | Chronically Ill with multiple symptoms | 4 | Age range: 44–63 |  | Phase 1 10 daily consecutive sessions. Phase 2; as needed over 20 days. | 20 Min | DLPFC montage (left anodal) or MI-SO electrode montage | Telehealth-tDCS protocol was successful and easy to replicate electrode placement at home via headband – pre-determined position. |
| Shaw et al. (2017) | Double Blind Randomized Controlled Trial | Multiple Sclerosis (MS) and Parkinson Disease (PD) | <u>Study 1</u><br>26 (MS)<br><u>Study 2</u><br>MS=20 & PD=6 | No Info. | <u>Study 1</u><br>=1.5<br><u>Study 2</u><br>MS=2<br>PD= 2 or 1.5 | <u>Study 1</u> =10<br><u>Study 2</u><br>MS=20<br>PD=10 | 20 Min | DLPFC (left anodal) | Total of 748 sessions completed with high tolerability. tDCS is feasibility with remote supervision. |

**Table 2:** Specific Elements of Transcranial Direct Current Stimulation Delivered Remotely.

| Authors (Year) | Pre Training on tDCS | Pre and/or Paired Intervention with tDCS | Home visit By Research team | Additional features or Equipment to support remote delivery | Monitoring/Support/fidelity | Remarks |
| --- | --- | --- | --- | --- | --- | --- |
| Agarwal et al. (2018) | 1<sup>st</sup> session: training for participants at clinic | Paired: Cognitive Training | No | -Home tDCS kit, laptop with a mouse & software<br>-Program for remote control & video conferencing | -Video conferencing & remote control<br>-tDCS only “unlocks” one dose per code – controlled by study technician<br>-Family member also trained on tDCS in case further assistance is needed | -All sessions were well tolerated and completed successfully<br>-The telemedicine protocol for PD patients maximized compliance & recruitment<br>-Afternoon sessions were more effective than morning sessions |
| Andrade (2013) | N/A | Medication; Clozapine (200-300 mg/d); Aripiprazole (15mg/d) | N/A | No | Medically qualified family member | -Domiciliary tDCS is a feasible treatment but needs to be monitored frequently to confirm adherence to treatment protocols<br>-No long-term harm |
|  |  | -Patient previously on rTMS treatment |  |  |  |  |
| Andre et al. (2016) | No info | No | No | No | No | No info |
| Bystad et al. (2017) | No info, but indicated patient understood the procedure | No | No info | -Fixed stimulation schedule (8 am) and reminder on patient's phone<br>-Stimulator output controlled with multimeter device | Patient's wife | Long term self-administration of tDCS for AD patient was possible with support from caregiver (wife). |
| Carvalho et al. (2018) | 1 training session: step by step process for self-stimulation | No | N/A | Phone to communicate with research team | -Diary<br>-Written form<br>-Contact phone (Team available 24 hr via phone) | Adherence was high (90%). tDCS is feasible for home use with monitoring. |
| Cha et al. (2016) | 3 days (30-60 min each) of tDCS self-administration training | Pre: 5 consecutive sessions of rTMS | No | Participants own cell phone | -Personalized web links through SurveyMonkey with a daily check in<br>-‘Study Buddy’ for back-up communication<br>-Email & phone calls | The home based self-administered tDCS was found to be excellent and very safe. Compliance was high and participant felt confident setting up tDCS. |
| Charvet et al. (2017) | Participant trained on tDCS, tolerability | Paired: Cognitive Training | No | Study kit at-home use (laptop computer with mouse and charger, tDCS device) | Supervised at all times during sessions via videoconferencing software. | Home based tDCS treatment is possible with RS-tDCS protocol among people with MS. |
|  | testing followed by 1<sup>st</sup> tDCS session at clinic |  |  | with headset, sponges, and extra saline) |  |  |
| Charvet et al. (2017) | 1<sup>st</sup> session at clinic to determine capacity for self-administration of tDCS | Paired: Cognitive Training | No | -Laptop with software for real time supervision.<br>-A one-time use dose code for each session. | -Videoconferencing and program for remote access by study technician.<br>-Informational packet & short instructional video.<br>-A proxy or caregiver to assist with tDCS headset placement and device operation | Successful in reaching participants away from clinic to conduct self-administrated tDCS via telerehabilitation protocols. |
| Hagenacker et al. (2014) | 1<sup>st</sup> session training for participants and relative at research centre. | Participants on stable medication and anti-epileptic drugs. | No | Phone (if needed) | Trained relative, diary, electronic protocol within stimulator that records correct tDCS application, phone (in case of problems; but never used by participant). | tDCS was successful in reducing pain through self-evaluation, but remote delivery was problematic for elderly patients and drop-out rates were high. |
| Hyvarinen et al. (2016) | 1<sup>st</sup> session completed at outpatient clinic after a training session. | No | No | Diary, free-form notes & instructions to monitor and report skin condition. | No supervision provided; Patients keep treatment diary, and all the tDCS parameters were pre-programmed into the tDCS device. | Self-administered tDCS was easy and safe; proper training with pre-screening for suitability is essential. Supervision from a healthcare professional is recommended. |
| Kasschau et al. (2015) | 1<sup>st</sup> 2 sessions used for in person training | No | Study technician visited during 2<sup>nd</sup> session to confirm | Laptop with instructional video & secure video conferencing | Web Conferencing | All 152 remotely supervised sessions showed 100% success in correctly placing electrodes and operating tDCS device. |
|  |  |  | correct set-up and assess home suitability | connection with technician. |  |  |
| Kasschau et al. (2016) | 1<sup>st</sup> session: training for participants (or proxy) at clinic | Paired: Web based adaptive cognitive training | Home visit during 2<sup>nd</sup> session for equipment delivery & to oversee the first virtual session. | -Laptop with telemedicine software<br>-Participant's cell phone at their workstation<br>-One-time use dose code to unlock each session, controlled by a study technician. | -Web conferencing<br>-Participants cell phone used for back up communication | -None of the sessions were discontinued for any participants<br>-100% complete adherence rate during the stimulation protocol. |
| Marten et al. (2018) | Training at home or nursing home for family member or caregiver only | No | Patient seen at home or nursing home for training | No | -tDCS device recorded sessions for adherence<br>-Patient's relative and/or caregiver gave daily report of any abnormalities | -All patients tolerated tDCS<br>-Home-based tDCS can be used outside a research facility or hospital by patients or caregivers. |
| Mortensen et al. (2015) | Yes, but no information available | Paired: Occupational Therapy | tDCS applied for remote training by occupational and physiotherapists | Delivered by trained occupational and physiotherapists at participants home | Supervised by primary investigator but no further information available | -tDCS can be easily applied for home-based rehabilitation by occupational/ physiotherapists following practical and theoretical instruction<br>-tDCS stimulation well tolerated by participants |
| Riggs et al. (2018) | One training session at home | No | One home visit for initial phase; eligibility, tolerability & training | Telehealth device paired with tDCS | -Telehealth device which allowed remote assistance, adherence monitoring, and videoconferencing<br>-Informal caregiver | -No difficulties with training participants, protocol adherence, or tolerability<br>-60 sessions completed without discontinuation or adverse events. |
| Shaw et al. (2017) | 1<sup>st</sup> session: trained patients at clinic as per RS-tDCS protocol. | Paired: Cognitive Training | No | Computer with the use of remote desktop software | Web conferencing | RS-tDCS protocol is feasible. |

**Table 3:** Summary Characteristics of Studies on Remotely-Delivered Cognitive Remediation and Cognitive Training.

| Authors (Year) | Disease | N/Group Condition | Age (SD) | Design | Number of Sessions/ Period | Setting (Individual/ Group/ Couple) | Outcome Measures | Results |
| --- | --- | --- | --- | --- | --- | --- | --- | --- |
| Anguera et al. (2017) | Sensory Processing Dysfunction (SPD) | <u>Experiment 1</u> = 62 (20 SPD +ADHD, 25 Control & 17 SPD only)<br><u>Experiment 2</u> = 57 (Final analysis: 17, 22 & 10) | 9.7 (1.3) SPD +ADHD, 10.5(1.3) Healthy Control, 10.3(1.5) SPD | Pilot Study; experimental design | 30 mins per day which consists of 7 tasks, 3 – 4 minutes sessions, 5 days per week for 4 weeks | Patient (a child) and their caregiver (parent) | Perceptual discrimination task, Test of Variables of Attention (TOVA) & EVO assessment (perceptual discrimination, visuomotor tracking and multitasking ability) | -SPD children with inattention/hyperactivity showed improvement in midline frontal theta activity and in inattention.<br>-Experiment 1 and 2 used the same participants, but Experiment 2 used remote cognitive training. |
| Boman et al. (2004) | Mild to moderate acquired non- | 10 | 47.5 | Pre-post-follow-up design | 1 hour, three times weekly for 3 weeks | Individual | The Attention Process Training test, Digit Span Test, Claeson-Dahl test, | Significant improvement in attention, |
|  | progressive brain injury |  |  | (single group) | in their home or at work |  | The Rivermead Behavioural Memory test, The Assessment of Motor and Process Skills, The European Brain Injury Questionnaire, Self-perceived quality of life. | concentration and memory.<br>-No significant improvement in activity. |
| Caller et al. (2016) | Epilepsy | 66 randomized to 3 equal groups. Final analysis: 15 in H, 14 in H+ (coupled with memory training) and 20 control | 49.3(9.2) H/H+ and 41.4(11.2) control | Randomized control trial | 20 – 40 min daily, 5 days a week for 8 weeks | Individual | Quality of Life in Epilepsy scale, QOLIE-31, RBANS, PHQ-9, FACT-Cog, BRIEF-A and Satisfaction Survey | Significant improvement in cognition and quality of life. |
| Charvet et al. (2015) | Multiple Sclerosis | 20 (11 Experiment & 9 Active Control) | 19 - 55 | Double blind randomized control trial | 30 min per day/5 days a week over 12 weeks (Target: 60 total days played across 3 months) | Individual | <u>Cognitive Composite:</u><br>-WAIS-IV (LNS), SRT, BVMT-R, Corsi block visual sequence<br><u>Motor Composite:</u><br>-DKEFS trials, Nine-hole peg test, Timed 25-foot walk<br><u>Self-report measures:</u><br>-ECog | Significant improvement in cognitive measures and motor tasks. |
| Charvet et al. (2017) | Multiple Sclerosis | 135 (74 Experiment & 61 Active control) | 50 (12) | Double blind randomized control trial | 1 hour per day, 5 days a week over 12 weeks (Total target: 60 hours) | Individual | Neuropsychological Test – PASAT, WAIS-IV (LNS & DSB), BVMT-R, D-KEFS, (2) Self Report change in Cognition. | Significant improvement in cognitive functioning. |
| Cody et al. (2015) | HIV | 20 | 50.22(6.57) | Within subjects pre-post experiment | 2 hours per week for 5 weeks (Target is 10 hours) | Individual | Useful Field of View (UFOV®) Wisconsin Card Sorting, Finger Tapping, Timed IADL measures and feedback on training | Significant improvement in processing speed and possible transfer to activity of daily living. |
| Fisher et al. (2009) | Schizophrenia | 55 (29 experiment auditory training & 26 control-computer games)<br>-Only 10 in exp. group performed remote training | <u>Experiment</u><br>42.86 (10.07)<br><u>Control</u><br>45.31(9.39) | Pre-post; controlled experiment design | 1 hour per day, 5 days per week for 10 weeks | Individual | PANSS, Quality of Life – Abbreviated Version and MATRICS | -Participants in experiment group showed significant improvement in cognition, memory and auditory psychophysical performance.<br>-Data analysis combined training from both remote & laboratory settings. |
| Fisher et al. (2015) | Schizophrenia | 86 (43 experiment & 43 Control group) | 21.22 | Double blind randomized control trial | 1 hour daily, 5 days per week for 8 weeks (40 hours training) | Individual | MATRICS, D-KEFS Tower Test, Strauss Carpenter Outcome, and Global Functioning: Role and Social Scales | Participants in experiment group (auditory training) showed significant improvement in cognition, memory and problem solving. |
| Johnstone et al. (2017) | ADHD | Total 107; 54 experiment (44 completed) & 53 control (41 completed) | N/A | Randomized waitlist control design | 25 sessions over a period of 6 to 8 weeks (3 or 4 sessions a week) | Patient (child) and their caregiver (parent) | CBCL, Conners 3-P, ADHD-RS, and WIAT-11 | Trainees improved in the trained tasks but enjoyment and engagement declined. |
| Kirk et al. (2016) | Intellectual and developmental disabilities (IDD) | 76 (38 Experiment & 38 Control; 37 in final analysis) | 8.22 | Double blind randomized control trial | About 20 min per day, 5 times per week, over a 5 week period | Patient (a child) and their caregiver (parent) | WATT and SWAN | Children that received home based attention training showed greater improvement in selective attention performance. |
| Loewy et al. (2016) | Clinical High Risk (CHR) patients for psychosis | 83 (Experiment 50; only 31 completed & Control 33; only 17 completed) | 18.1 | Double blind randomized control trial | 1 hour per day, 5 days a week for 8 weeks (40 hours total) | Individual | SOPS, Global Functioning: Role and Social Scales, MATRICS, D-KEFS, NAB Mazes, HVLT-R and BVMT-R | Participants in experiment group showed significant improvement in verbal memory. |
| Mariano et al. (2015) | 22q11 Deletion Syndrome | Enrolled: 22<br>Final analysis: 21 | 14.6 (1.3) | Longitudinal within-group design | 45 min per day, 3 times per week for 8 months | Individual (teleconference) | Neurocognitive test battery; CNS Vital Signs (CNS-VS) | Significant improvement in working memory, shifting attention and cognitive flexibility. |
| McBride et al. (2017) | Chronic Fatigue Syndrome (CFS) | 76 (36 CBT/GET program & 36 CBT/GET + CR program) | 35.5 (age range: 13 – 71) | Case control trial | 3-5 sessions per week, up to a total of 40 sessions | Individual | SPHERE (SOMA & PSYCH subscale), SF-36, Neuropsychological Performance measures | Significant improvement in neurocognitive symptoms and cognition. |
| Milman et al. (2014) | Parkinson's Disease | 18 | 67.7 (6.4) | Pre-post; single group experiment | 30 min a day, 3 days per week for 12 weeks | Individual | The Mindstreams (NeuroTrax Corp., TX) battery of computerized neuropsychological test and the Timed Up and Go (TUG) test | Significant improvement in global cognitive score & Timed Up and Go (TUG) measures. |
| Mohanty & Gupta (2013) | Traumatic Brain Injury (TBI) | 1 | 24 | Single case study | 45 min to 1 hour twice a day, for 9 months | Individual and parent (father) | PGI Battery of Brain Dysfunction, Selected tests from NIMHANS & Dysfunctional Analysis Questionnaire | Improvement in cognitive functions and day to day functioning. |
| Nahum et al. (2014) | Schizophrenia | 34 (17 Schizophrenia & 17 matched healthy control) | 23.7 | Pilot experimental study & within subject design | 1-2 hours per day, 2–5 days per week for 6-12 weeks (24 hours) | Individual | SocialVille Training Program Feasibility and ease of use, SocialVille Exercise-based Assessments, Penn Facial Memory, PROID, MSCEIT), Social and Role Scales, SFS, QLS, BIS/BAS, TEPS | Improvement on speeded SocialVille and working memory tasks, motivation, social cognition and functioning.<br>-Only 10 Schizophrenia patients engaged in cognitive training from home. |
| Pyun et al. (2009) | Stroke (Cognitive Impairment) | Recruited 6 (2 did not complete the full 12 week home program) | 48.7 (age range 28 – 62) | Multiple case study | 2 hours (only 30 min of CR) per day, 7 days a week for 12 weeks | Patients and their caregivers | MMSE, NCSE, domain-specific computerized neuropsychological test, LOTCA, MBI & S-IADL | Significant improvement in activity of daily living and marginal improvement in general cognition. |
| Quayhagen et al. (2001) | Alzheimer's Dementia | <u>Experiment 1</u><br>56 couples (experimental vs. placebo vs. control)<br><u>Experiment 2</u><br>30 couples (experimental vs. control) | <u>Experiment 1</u><br>Patients; 73.18 & caregivers 67.75<br><u>Experiment 2</u><br>Patients; 74.97 & caregiver 72.57 | Randomized control trial | <u>Experiment 1</u><br>1 hour a day, 5 days a week for 12 weeks.<br><u>Experiment 2</u><br>1 hour a day, 5 days a week for 8 weeks. | Spousal-caregiving units (patient and caregiver) | WMS-R, DRS, FAS, GCS & Marital Needs Satisfaction Scale. | -Improvement in immediate memory for experiment 1 and problem solving for experiment 2.<br>-Verbal fluency improved in both studies. |
| Quayhagen et al. (1995) | Alzheimer's Dementia | 79 patients and caregivers (78 in final analysis) | 73.6 (8.0) patients & 66.7 (10.8) caregivers | Randomized control trial | 1 hour a day, 6 days a week for 12 weeks | Patient and their caregiver | DRS, WMS-R, FAS, Geriatric Coping Schedule, Memory and Behavior Problems Checklist (part A) | Experiment group showed improvement in cognition and behavioral performance. |
| Rajeswaran et al. (2017) | Schizophrenia | 1 | 26 | Pre-post intervention single case study | 1 hour a day for 10 weeks | Individual (patient) and caregiver (mother) | NIMHANS & social functioning | Cognitive retraining improved cognitive functions. |
| Regan et al. (2017) | Mild Cognitive Impairment (MCI) & Alzheimer's Dementia (AD) | 55 enrolled; 40 finished study (25 intervention & 15 control) | Client: 77.2 (6.5), Caregiver: 66.8 (15.0) | Multicenter randomized control trial | 1 hour per week for 4 weeks | Individual with their caregiver | COPM, HADS, ICQ, MMCQ, QOD, B-ADL, ECOQ, RMBPC | Intervention group showed significant improvement in performance and satisfaction. |
| Shaw et al. (2017) | Multiple Sclerosis (MS) and Parkinson Disease (PD) | <u>Study 1:</u> 26 (MS)<br><u>Study 2:</u> 20 (MS) & 6 (PD) | N/A | <u>Study 1:</u> Open label<br><u>Study 2:</u> Double blind randomized sham control study (PD arm open label) | Study 1: 20 min per day, 5 days a week for two weeks (9 session at home)<br>Study 2=20 & PD=10 | Individual | Feasibility report, UPDRS, NSNQ, PROMIS & PANAS | -Supports the feasibility of remotely administered tDCS paired with cognitive training.<br>-Study was ongoing at time of publication. |
| Vazquez-Campo et al. (2016) | Schizophrenia | 21 (12 intervention & 9 control); final = analysis 19 | 39.28 | Pre/post pilot study | 1 hour per week for 12 weeks | Individual | EP; Ekman 60 Faces Test, ToM; Hinting Task, Recognition of Faux Pas, Strange Stories of Happe, AIHQ; Ambiguous Intentions Hostility Questionnaire, MSCEIT, PANSS, WAIS-IV & Semi-structured interview | Significant improvement in EP, ToM and AS variables.<br>-Only 30% took part in the intervention from their home. |
| Ventura et al. (2013) | Schizophrenia | Recruited 9 (8 completed study) | N/A | Feasibility study | 1 hour twice per week for 6 weeks | Individual and relative | -MCCB composite, - CGI-Cogs – patient, informant (relative) and rater version, BQKC total score – patient & relative version, SCORS social & work functioning | Improvement in cognition, knowledge (about the role of cognition in daily life), and improvement in social functioning. |

**Table 4:** Specific Elements of Intervention Using Remotely-Delivered Cognitive Remediation and Cognitive Training.

| **Authors (Year)** | **Software Name** | **Pre-Training at Lab/Study site** | **Intervention Name/CR Program/Content** | **Mode of Delivery** | **Training/Monitoring/Support/Fidelity** | **Remark** |
| --- | --- | --- | --- | --- | --- | --- |
| Anguera et al. (2017) | EVO by Akili Interactive Labs | Self-guided treatment | Cognitive training involves a combination of visuomotor and perceptual discrimination tasks | iPad with internet | -Research assistant provided remote monitoring with support and feedback during training as well as reminder phone calls. Parent also available for support<br>-EVO gives real-time feedback and adaptive algorithms | Highlights the benefit of targeted attention intervention via EVO (both assessment & intervention) at their own home for children with SPD. |
| Boman et al. (2004) | N/A | N/A | 1-APT-training 2-Generalization 3-Teaching of compensatory strategies for self-selected cognitive problems | Computer | N/A | Home based cognitive training improved cognition and supported the learning of strategies. |
| Caller et al. (2016) | Home Based Self-Management and Cognitive Training Changes lives (HOBSCOTCH), Nintendo DS® & Brain Age©program | First session in a group format with introduction to their 'memory coach' | -HOBSCOTCH; Self-efficacy principles combined with compensatory strategies to optimize functioning<br>-Computerized memory training | Handbook & worksheet, computer & device for games | -Sessions conducted over the phone by epilepsy specialized ARNP or RN trained as 'memory coaches'<br>-Device to record completion of exercises to allow for compliance monitoring | HOBSCOTCH could be a good cognitive remediation program option for patients with epilepsy who experience transportation barriers. Patients reported high satisfaction rate and over 70% preferred the 'telephone visit' |
| Charvet et al. (2015) | Lumos Labs Inc., "TeamViewer" for remote support and "WorkTime" software by NesterSoft Inc. for time | N/A | -Adaptive cognitive training program using Lumosity platform<br>-Hoyle puzzles and board game program for active control | 17" laptop computer with internet, noise cancelling headsets, hand-held mice. Wifi provided if | Technical support, coaching, and monitoring of computer use were provided remotely by a study technician | -High compliance rate<br>-Potential meaningful benefit for person with MS<br>-Able to reach participants away from clinic setting, and facilitated rapid recruitment at a much lower cost. |
|  | tracking and monitoring. |  |  | no internet available |  |  |
| Charvet et al. (2017) | Brain HQ program (CR training) & “Work Time” (monitor & record real time) | No prior training provided | Telerehabilitation; Adaptive Cognitive Remediation (ACR). | 17” laptop computer with internet, headphones, and a user guide | -Ongoing access to technical support, a monitoring software program and scheduled weekly check in phone calls by an unblinded study technician | Telerehabilitation approach allowed rapid recruitment and high compliance rate. 135 participants were recruited within 12 months of trial. |
| Cody et al. (2015) | RoadTour by POSIT Science | Participants given software (CD) and written instruction on how to install the program. | Processing speed tasks used a double-stair case technique | Personal computer | -Participants able to call study staff for support<br>-Participants kept a log to record training activities | Home-based computerized cognitive training program can be administered remotely among people with HIV to improve processing speed. |
| Fisher et al. (2009) | Computerized software exercises (no name given) | N/A | Auditory training exercises | Computer | -Weekly call by research staff<br>-Compliance monitored through electronic data following each training session | Total mean training time was 47.9 hours (SD=7.5). There was no separate data for participants who completed remote training. |
| Fisher et al. (2015) | Posit Science | Coaching was provided if participant was having difficulties completing recommended number of hours/week | -Computerized Auditory Training (AT) to improve speed and information processing<br>-Control group played commercially available games | Loaned laptop computer with internet | -Coaching if necessary<br>-Compliance monitored by electronic data upload<br>-Phone contact 1-2 times a week to discuss progress. | 40 participants from each condition completed 20–40 hours of auditory training and computer games respectively. |
| Johnstone et al. (2017) | Focus Pocus by Neurocognitive Solutions Pty Ltd. | Pre-training demonstration with participants & their parent(s) | Cognitive training combined with neuro EEG feedback. Each session consisted of 14 mini-games | Participant’s computer | EEG was recorded continuously from site Fp1 at 256 Hz. | Technology development supports intervention to be deliverable at home; Neurofeedback training can reduce ADHD symptoms. |
| Kirk et al. (2016) | The Training Attention and Learning Initiative (TALI) | Initial session at University (research centre) and school | Computerized program targets attention skills via four activities e.g. fish tank | 7” touch screen tablet | -Supervision at home by parent/guardian. Research assistant contacted participants to monitor progress and to give technical support | High compliance rate: 34/38 (90%) participants in cognitive training met compliance criteria. |
|  |  |  |  |  | -Reward system used |  |
| Loewy et al. (2016) | Posit Science | N/A | Auditory processing-based exercises, verbal learning & memory operations | Loaned laptop computer | -Participants were contacted 1–2 times per week via phone to monitor progress. Phone survey completed after 10 hours to monitor adherence<br>-Coaching provided for participants facing difficulties in completing task. Point rewards system used and compliance monitored by electronic data | -CHR individuals benefited from home-based cognitive training.<br>-Hours of training = 21.5 (16.3) for both groups.<br>-High study attrition rate (42%). |
| Mariano et al. (2015) | Adapted from Computerized CogRehab system | No prior training provided | Challenging Our Mind (COM) | Laptop computer with built-in camera, internet & Cisco WebEx web conferencing | -CR intervention conducted by “Cognitive Coach” via Web conference. Coaches had biweekly meetings with PI for monitoring. COM sessions were recorded & reviewed by independent rater<br>-Instructional Manual Strategies Guide (available by request) | -Youth with 22q11DS successfully connected from their own home.<br>-Increased service provision in rural settings. |
| McBride et al. (2017) | Online CR Training Program by Lumos Labs Inc. | Initially with CBT and GET sessions | -CR combined with CBT and GET.<br>-CR had 24 different game-based tasks to train attention, working memory, processing speed and executive functioning | Personal computer with internet | -Program gives automatic visual and auditory feedback as well as reinforcement regarding performance<br>-Guidelines, frequency of sessions and breaks within the session were set by clinical psychologist. Written and animated instructions also provided | Home-based cognitive remediation training program can be an effective intervention for people with CFS. |
| Milman et al. (2014) | Attengo® software (Attenfocus®) | N/A | Executive function and attention training; games involved problem solving, information processing etc. | Personal computer with internet | The system recorded the time spent on the exercises using software, and automatically | -Computerized cognitive training has a therapeutic benefit for people with PD. |
|  |  |  |  |  | sent it to the research team via internet | -Compliance rate was about 4 hours less than the recommended training time over the 12 weeks period. |
| Mohanty & Gupta (2013) | Home based cognitive retraining program | Task selected for a particular week was first demonstrated, then rehearsed by the co-therapist (father) | Neuropsychological remediation tasks in a graded fashion. There was also counselling and psychoeducation sessions to deal with anxiety and help with realistic expectation setting | N/A (likely printed materials) | -Patient's father as co-therapist<br>-Progress review & counselling once a week for the first 2 months, biweekly for the next 2 months and once a month for the last 6 months (18 sessions total). | Home based neuropsychological remediation program was found to be therapeutic in brain damaged patient. |
| Nahum et al. (2014) | SocialVille online program | N/A | Social cognition training intervention had 19 computerized exercises targeting speed, accuracy and processing of social information | Loaned laptop computers with internet | -Weekly phone calls and/or emails to monitor training adherence/requirement<br>-Clinical staff engaged in solution-focused conversations | -High adherence to online social cognition training.<br>-Reported medium to high satisfaction, enjoyment, and ease of use. |
| Pyun et al. (2009) | Individualized cognitive remediation with structured educational materials | Patient and their caregiver received a printed sheet of instructions and patient-recommended strategies with explanations | -Training material consisted of multi-level tasks to enhance attention, memory and executive function<br>-Content included CR, storytelling, recreational cognitive games, and aerobic exercise | -Printed material | -Checklist for caregivers to evaluate the patients' performance daily<br>-Weekly meeting with occupational therapist to answer questions, problems, monitoring on performance and progress rate. Program adjusted if needed. | Individualized home program found to be beneficial for chronic stroke patients with cognitive impairment. |
| Quayhagen et al. (1995) | Active cognitive stimulation material | Caregiver and patient trained in program. Return demonstrations by caregivers were required to validate training. | An instruction workbook for families with 12 modules to stimulate memory, problem solving and social interaction. | Printed workbook/ worksheets | Caregiver gave positive feedback, completed a weekly log book on progress, problems & successes. | This study supports the implementation of cognitive training at home among people with dementia. |
| Quayhagen et al. (2001) | Active cognitive stimulation material (Quayhagen & Quayhagen, 1989) | Caregiver learned from research team by observation and modelling | Materials to stimulate memory, fluency, and problem solving activities. | Printed materials | -Caregiver<br>-Research team provided 1 hour of weekly instruction to the patient at home. | Active involvement of spousal caregivers was beneficial in implementing cognitive remediation at home for persons with dementia. |
| Rajeswaran et al. (2017) | N/A | Visited hospital once a week | Cognitive retraining program by Hegde et al. (2008) plus a family intervention | N/A | N/A | -Patient also received EEG NFT for 40min/20 sessions.<br>-Patient and mother reported improvement. |
| Regan et al. (2017) | N/A | No | MAXCOG | Experienced counsellor at home | Counsellor and primary caregiver | MAXCOM is a brief but effective cognitive intervention. |
| Shaw et al. (2017) | Brain HQ (CR) & Team Viewer (real time monitoring) | First session involved training for participants at the clinic | Cognitive training targeted working memory, attention, processing speed etc. | Computer with remote desktop software | -Computers enabled for real-time monitoring and remote control by study staff<br>-Researcher monitored via Video-conferencing | High compliance rate and a successful protocol developed for remote tDCS + cognitive training self-administration. |
| Vazquez-Campo et al. (2016) | e-Motional Training (ET) | N/A | ET Training modules on emotional perception and a short animated cartoon | Computer and Internet | -Researcher monitored the patients' progress and resolved questions regarding computer and software<br>-Automatic metacognitive feedback with strategies | -Supports the feasibility of an online intervention to improve social cognition.<br>-Most participants found ET to be easy, entertaining and useful. |
| Ventura et al. (2013) | Posit Science | Each person with schizophrenia and their relative received 2 hours of in-office training on PositScience | Internet-based brain fitness program; targets critical cognitive functions using auditory discrimination tasks | Personal computer and internet | -Relative played an active role in software installation, planning, monitoring & emotional support for patient<br>-Regular phone contact with research team | -80% adherence to cognitive training at home.<br>-Supports feasibility using home-based cognitive training for people with Schizophrenia. |

## REFERENCES

1. Organization, W. H. (2017). The Top 10 Causes of Death. [cited 2017 December 12]; Available from: http://www.who.int/mediacentre/factsheets/fs310/en/.

2. Shaw, M. T., Kasschau, M., Dobbs, B., Pawlak, N., Pau, W., Sherman, K., Bikson, M., Datta, A., & Charvet, L. E. (2017). Remotely Supervised Transcranial Direct Current Stimulation: An Update on Safety and Tolerability. Journal of Visualized Experiments,(128), 1–8. DOI: 10.3791/56211.

3. Organization, W. H. (2017). Dementia. [cited 2017 December 12]; Available from: http://www.who.int/mediacentre/factsheets/fs362/en/.

4. Bowie, C. R., Gupta, M., Holshausen, K., Jokic, R., Best, M., & Milev, R. (2013). Cognitive remediation for treatment resistant depression: Effects on cognition and functioning and the role of online homework. Journal of Nervous and Mental Disorders, 201(8), 680–685. DOI: 10.1097/NMD.0b013e31829c5030.

5. Kuo, M., Paulus, W., & Nitsche, M. A. (2014). Therapeutic effects of non-invasive brain stimulation with direct currents (tDCS) in neuropsychiatric diseases. NeuroImage, 85(3), 948–960. DOI: https://doi.org/10.1016/j.neuroimage.2013.05.117.

6. Rajji, T. K. (2019). Impaired brain plasticity as a potential therapeutic target for treatment and prevention of dementia. Expert Opinion on Therapeutic Targets, 23(1), 21–28. DOI: https://doi.org/10.1080/14728222.2019.1550074.

7. Ferrucci, R., Mameli, F., Guidi, I., Mrakic-Sposta, S., Vergari, M., Marceglia, S., Cogiamanian, F., Barbieri, S. Scarpini, E. & Priori, A. (2008). Transcranial direct current stimulation improves recognition memory in Alzheimer disease. Neurology, 71(7), 493–498. DOI: https://doi.org/10.1212/01.wnl.0000317060.43722.a3.

8. Naismith, S. L., Redoblado-Hodge, M. A., Lewis, S. J., Scott, E. M., & Hickie, I. B. (2010). Cognitive training in affective disorders improves memory: A preliminary study using the NEAR approach. Journal of Affective Disorders, 121(3), 258–262. DOI: doi:10.1016/j.jad.2009.06.028.

9. Boggio, P. S., Khoury, L. P., Martins, D. C., Martins, O. E., Macedo, E. C., & Fregni, F. (2009). Temporal cortex direct current stimulation enhances performance on a visual recognition memory task in Alzheimer disease. Journal of Neurology, Neurosurgery & Psychiatry, 80(4), 444–447. DOI: doi:10.1136/jnnp.2007.141853.

10. Boggio, P. S., Ferrucci R., Mameli F., Martins D., Martins O., Vergari M., Tadini L., Scarpini E., Fregni F., & Priori A. (2012). Prolonged visual memory enhancement after direct current stimulation in Alzheimers disease. Brain Stimulation, 5(3), 223–230. DOI: doi:10.1016/j.brs.2011.06.006.

11. Grill, J. D., & Karlawish, J. (2010). Addressing the challenges to successful recruitment and retention in Alzheimers disease clinical trials. Alzheimers Research & Therapy, 2(6), 34–44. DOI: doi:10.1186/alzrt58.

12. Deckersbach, T., Nierenberg, A. A., Kessler, R., Lund, H. G., Ametrano, R. M., Sachs, G., Rauch, S.L., & Dougherty, D. (2010). Cognitive Rehabilitation for Bipolar Disorder: An Open Trialfor Employed Patients with Residual Depressive Symptoms. CNS Neuroscience & Therapeutics, 16(5), 298–307. DOI: doi: 10.1111/j.1755-5949.2009.00110.x.

13. Meusel, L. A., McKinnon, M., Hall, G., & MacQueen, G. M. (2009). Evidence for sustained improvement in memory deficits following computer-assisted cognitive remediation in patients with a mood disorder. Biological Psychiatry, 65(8).

14. Rupp, C. I., Kemmler, G., Kurz, M., Hinterhuber, H., & Fleischhacker, W. W. (2012). Cognitive Remediation Therapy During Treatment for Alcohol Dependence. Journal of Studies on Alcohol and Drugs, 73(4), 625–634. DOI: doi:10.15288/jsad.2012.73.625.

15. Wykes, T., Huddy, V., Cellard, C., McGurk, S. R., & Czobor, P. (2011). A meta-analysis of cognitive remediation for schizophrenia: Methodology and effect sizes. American Journal of Psychiatry, 168(5), 472–485.

16. Bowie, C. R., 2016 Cognitive Remediation for Major Depressive Disorder, in Cognitive impairment in major depressive disorder: Clinical relevance, biological substrates, and treatment opportunities, R. S. C. McIntyre, D. S., Editor. Cambridge University Press: Cambridge.

17. Brunoni, A. R., Nitsche, M. A., Bolognini, N., Bikson, M., Wagner, T., Merabet, L., Edwards D., J., Valero-Cabrei, A., Rotenberg A., Pascual-Leone, A., Ferruccil, R., Priori, A., Boggio, P.S., & Fregni, F. (2012). Clinical research with transcranial direct current stimulation (tDCS): Challenges and future directions. Brain Stimulation, 5(3), 175–195. DOI: doi:10.1016/j.brs.2011.03.002.

18. Hagenacker, T., Bude, V., Naegel, S., Holle, D., Katsarava, Z., Diener, H. C., & Obermann, M. (2014). Patient-conducted anodal transcranial direct current stimulation of the motor cortex alleviates pain in trigeminal neuralgia. The Journal of Headache and Pain, 15 78. DOI: https://doi.org/10.1186/1129-2377-15-78.

19. Charvet, L. E., Shaw, M. T., Haider, L., Melville, P., & Krupp, L. B. (2015). Remotely-delivered cognitive remediation in multiple sclerosis (MS): protocol and results from a pilot study. Mult Scler J Exp Transl Clin, 1 2055217315609629. DOI: 10.1177/2055217315609629.

20. Kasschau, M., Reisner, J., Sherman, K., Bikson, M., Datta, A., & Charvet, L. E. (2016). Transcranial direct current stimulation is feasible for remotely supervised home delivery in multiple sclerosis. Neuromodulation.

21. Quayhagen, M. P., Quayhagen, M., Corbeil, R. R., Roth, P. A., & Rodgers, J. A. (1995). A dyadic remediation program for care recipients with dementia. Nursing Research, 44(3), 153–159. DOI: http://dx.doi.org/10.1097/00006199-199505000-00005.

22. Quayhagen, M. P., & Quayhagen, M. (2001). Testing of a cognitive stimulation intervention for dementia caregiving dyads. Neuropsychological Rehabilitation, 11(3-4), 319–332. DOI: https://doi.org/10.1080/09602010042000024

23. Regan, B., Wells, Y., Farrow, M., Ohalloran, P., & Workman, B. (2017). MAXCOG— Maximizing Cognition: A Randomized Controlled Trial of the Efficacy of Goal-Oriented Cognitive Rehabilitation for People with Mild Cognitive Impairment and Early Alzheimer Disease. The American Journal of Geriatric Psychiatry, 25(3), 258–269. DOI: 10.1016/j.jagp.2016.11.008.

24. Bystad, M., Rasmussen, I. D., Gronli, O., & Aslaksen, P. M. (2017). Can 8 months of daily tDCS application slow the cognitive decline in Alzheimer’s disease? A case study. Neurocase, 23(2), 146–148. DOI: https://doi.org/10.1080/13554794.2017.1325911.

25. Andre, S., Heinrich, S., Kayser, F., Menzler, K., Kesselring, J., Khader, P. H., Lefaucheur, J. P., & Mylius, V. (2016). At-home tDCS of the left dorsolateral prefrontal cortex improves visual short-term memory in mild vascular dementia. Journal of the Neurological Sciences, 369 185–190. DOI: https://doi.org/10.1016/j.jns.2016.07.065.

26. Charvet, L., Shaw, M., Dobbs, B., Frontario, A., Sherman, K., Bikson, M., Datta, A., Krupp, L., Zeinapour, E., & Kasschau, M. (2017). Remotely Supervised Transcranial Direct Current Stimulation Increases the Benefit of At-Home Cognitive Training in Multiple Sclerosis. Neuromodulation. DOI: 10.1111/ner.12583.

27. Charvet, L. E., Dobbs, B., Shaw, M. T., Bikson, M., Datta, A., & Krupp, L. B. (2017). Remotely supervised transcranial direct current stimulation for the treatment of fatigue in multiple sclerosis: Results from a randomized, sham-controlled trial. Multiple Sclerosis, 1352458517732842. DOI: 10.1177/1352458517732842.

28. Kasschau, M., Sherman, K., Haider, L., Frontario, A., Shaw, M., Datta, A., Bikson, M., & Charvet, L. (2015). A Protocol for the Use of Remotely-Supervised Transcranial Direct Current Stimulation (tDCS) in Multiple Sclerosis (MS). Journal of Visualized Experiments,(106), 1–8. DOI: 10.3791/53542.

29. Andrade, C. (2013). Once-to Twice-Daily, 3-Year Domiciliary Maintenance Transcranial Direct Current Stimulation for Severe, Disabling, Clozapine-Refractory Continuous Auditory Hallucinations in Schizophrenia. Journal of ECT, 29(3), 239–242. DOI: 10.1097/YCT.0b013e3182843866.

30. Agarwal, S., Pawlak, N., Cucca, A., Sharma, K., Dobbs, B., Shaw, M., Charvet, L., & Biagioni, M. (2018). Remotely-supervised transcranial direct current stimulation paired with cognitive training in Parkinson’s disease: An open-label study. Journal of Clinical Neuroscience, 57 51–57. DOI: 10.1016/j.jocn.2018.08.037.

31. Mortensen, J., Figlewski, K., & Andersen, H. (2016). Combined transcranial direct current stimulation and home-based occupational therapy for upper limb motor impairment following intracerebral hemorrhage: a double-blind randomized controlled trial. Disability and Rehabilitation, 38(7), 637–643. DOI: https://doi.org/10.3109/09638288.2015.1055379.

32. Hyvärinen, P., Mäkitie, A., & Aarnisalo, A. A. (2016). Self-Administered Domiciliary tDCS Treatment for Tinnitus: A Double-Blind Sham-Controlled Study. PLoS ONE 11(4). DOI: https://doi.org/10.1371/journal.pone.0154286

33. Carvalho, F., Brietzke, A. P., Gasparin, A., dos Santos, F. P., Vercelino, R., Ballester, R. F., Sanches, P. R., da Silva Jr, D. P., Torres, I. L., Fregni, F., & Caumo, W. (2018). Home-Based Transcranial Direct Current Stimulation Device Development: An Updated Protocol Used at Home in Healthy Subjects and Fibromyalgia Patients. Journal of Visualized Experiments,(137), 1–9. DOI: 10.3791/57614.

34. Cha, Y. H., Urbano, D., & Pariseau, N. (2016). Randomized Single Blind Sham Controlled Trial of Adjunctive Home-Based tDCS after rTMS for Mal De Debarquement Syndrome: Safety, Efficacy, and Participant Satisfaction Assessment. Brain Stimulation, 9(4), 537–544. DOI: https://doi.org/10.1016/j.brs.2016.03.016.

35. Martens, G., Lejeune, N., Obrien, A. T., Fregni, F., Martial, C., Wannez, S., Laureys, S., & Thibaut, A. (2018). Randomized controlled trial of home-based 4-week tDCS in chronic minimally conscious state. Brain Stimulation, 11(5), 982–990. DOI: 10.1016/j.brs.2018.04.021.

36. Riggs, A., Patel, V., Paneri, B., Portenoy, R. K., Bikson, M., & Knotkova, H. (2018). At-Home Transcranial Direct Current Stimulation (tDCS) With Telehealth Support for Symptom Control in Chronically-Ill Patients With Multiple Symptoms. Frontiers in Behavioral Neuroscience, 12(93), 1–10. DOI: 10.3389/fnbeh.2018.00093.

37. Ventura, J., Wilson, S. A., Wood, R. C., & Hellemann, G. S. (2013). Cognitive training at home in schizophrenia is feasible. Schizophr Res, 143(2-3), 397–8. DOI: 10.1016/j.schres.2012.11.033.

38. Mariano, M. A., Tang, K., Kurtz, M., & Kates, W. R. (2015). Cognitive remediation for adolescents with 22q11 deletion syndrome (22q11DS): a preliminary study examining effectiveness, feasibility, and fidelity of a hybrid strategy, remote and computer-based intervention. Schizophrenia Research, 166(1-3), 283–289. DOI: https://doi.org/10.1016/j.schres.2015.05.030.

39. Loewy, R., Fisher, M., Schlosser, D. A., Biagianti, B., Stuart, B., Mathalon, D. H., & Vinogradov, S. (2016). Intensive Auditory Cognitive Training Improves Verbal Memory in Adolescents and Young Adults at Clinical High Risk for Psychosis. Schizophrenia Bulletin, 42(suppl 1), S118–S126. DOI: https://doi.org/10.1093/schbul/sbw009.

40. Kirk, H. E., Gray, K. M., Ellis, K., Taffe, J., & Cornish, K. M. (2016). Computerised attention training for children with intellectual and developmental disabilities: a randomised controlled trial. Journal of Child Psychology and Psychiatry, 57(12), 1380–1389. DOI: https://doi.org/10.1111/jcpp.12615.

41. Anguera, J. A., Brandes-Aitken, A. N., Antovich, A. D., Rolle, C. E., Desai, S. S., & Marco, E. J. (2017). A pilot study to determine the feasibility of enhancing cognitive abilities in children with sensory processing dysfunction. PLoS One, 12(4), 1–19. DOI: https://doi.org/10.1371/journal.pone.0172616.

42. Caller, T. A., Ferguson, R. J., Roth, R. M., Secore, K. L., Alexandre, F. P., Zhao, W., Tosteson, T. D., Henegan, P. L., Birney, K., & Jobst, B. C. (2016). A cognitive behavioral intervention (HOBSCOTCH) improves quality of life and attention in epilepsy Epilepsy and Behavior, 57(Pt A), 111–117. DOI: https://doi.org/10.1016/j.yebeh.2016.01.024.

43. Milman, U., Atias, H., Weiss, A., Mirelman, A., & Hausdorff, J. M. (2014). Can cognitive remediation improve mobility in patients with Parkinson’s disease? Findings from a 12 week pilot study. Journal of Parkinson’s Disease, 4(1), 37–44. DOI: 10.3233/JPD-130321.

44. Mohanty, M., & Gupta, S. K. (2013). Home based neuropsychological rehabilitation in severe traumatic brain injury: a case report. Annals of Neurosciences, 20(1), 31–35. DOI: 10.5214/ans.0972.7531.200111.

45. Charvet, L. E., Yang, J., Shaw, M. T., Sherman, K., Haider, L., Xu, J., & Krupp, L. B. (2017). Cognitive function in multiple sclerosis improves with telerehabilitation: Results from a randomized controlled trial. PLoS One, 12(5), e0177177. DOI: 10.1371/journal.pone.0177177.

46. Fisher, M., Loewy, R., Carter, C., Lee, A., Ragland, J. D., Niendam, T., Schlosser, D., Pham, L., Miskovich, T., & Vinogradov, S. (2015). Neuroplasticity-based auditory training via laptop computer improves cognition in young individuals with recent onset schizophrenia. Schizophrenia Bulletin, 41(1), 250–258. DOI: https://doi.org/10.1093/schbul/sbt232.

47. Johnstone, S. J., Roodenrys, S. J., Johnson, K., Bonfield, R., & Bennett, S. J. (2017). Game-based combined cognitive and neurofeedback training using Focus Pocus reduces symptom severity in children with diagnosed AD/HD and subclinical AD/HD. International Journal of Psychophysiology, 116 32–44. DOI: 10.1016/j.ijpsycho.2017.02.015.

48. Vazquez-Campo, M., Marono, Y., Lahera, G., Mateos, R., & Garcia-Caballero, A. (2016). e-Motional Training(R): Pilot study on a novel online training program on social cognition for patients with schizophrenia. Schizophrenia Research: Cognition, 4 10–17. DOI: https://doi.org/10.1016/j.scog.2015.11.007.

49. Nahum, M., Fisher, M., Loewy, R., Poelke, G., Ventura, J., Nuechterlein, K. H., Hooker, C. I., Green, M. F., Merzenich, M., & Vinogradov, S. (2014). A novel, online social cognitive training program for young adults with schizophrenia: A pilot study. Schizophrenia Research: Cognition, 1(1), e11–e19. DOI: https://doi.org/10.1016/j.scog.2014.01.003.

50. Cody, S. L., Fazeli, P. L., & Vance, D. E. (2015). Feasibility of a Home-Based Speed of Processing Training Program in Middle-Aged and Older Adults With HIV. Journal of Neuroscience Nursing, 47(4), 247–254. DOI: 10.1097/JNN.0000000000000147.

51. Fisher, M., Holland, C., Merzenich, M. M., & Vinogradov, S. (2009). Using neuroplasticity-based auditory training to improve verbal memory in schizophrenia. The American Journal of Psychiatry, 166(7), 805–811. DOI: https://doi.org/10.1176/appi.ajp.2009.08050757.

52. Boman, I. L., Lindstedt, M., Hemmingsson, H., & Bartfai, A. (2004). Cognitive training in home environment. Brain Inj, 18(10), 985–95. DOI: 10.1080/02699050410001672396.

53. McBride, R. L., Horsfield, S., Sandler, C. X., Cassar, J., Casson, S., Cvejic, E., Vollmer-Conna, U., & Lloyd, A. R. (2017). Cognitive remediation training improves performance in patients with chronic fatigue syndrome. Psychiatry Research, 257 400–405. DOI: https://doi.org/10.1016/j.psychres.2017.08.035.

54. Pyun, S. B., Yang, H., Lee, S., Yook, J., Kwon, J., & Byun, E. M. (2009). A home programme for patients with cognitive dysfunction: a pilot study. Brain Injury, 23(7), 686–692. DOI: https://doi.org/10.1080/02699050902997862.

55. Rajeswaran, J., Taksal, A., & Jain, S. (2017). Rehabilitation in schizophrenia: A brain-behavior and psychosocial perspective. Indian Journal of Psychological Medicine, 39(6), 797–798. DOI: 10.4103/0253-7176.219648.

56. Charvet, L. E., Kasschau, M., Datta, A., Knotkova, H., Stevens, M. C., Alonzo, A., Loo C., Krull K.R., & Bikson, M. (2015). Remotely-supervised transcranial direct current stimulation (tDCS) for clinical trials: guidelines for technology and protocols. Frontiers in Systems Neuroscience, 9(26). DOI: 10.3389/fnsys.2015.00026.

57. Elgamal, S., Mckinnon, M. C., Ramakrishnan, K., Joffe, R. T., & Macqueen, G. (2007). Successful computer-assisted cognitive remediation therapy in patients with unipolar depression: a proof of principle study. Psychological Medicine, 37(9), 1229–1238. DOI: https://doi.org/10.1017/S0033291707001110.

